# TopFlash transgenic quail reveals dynamic TCF/β-catenin signaling during avian embryonic development

**DOI:** 10.1101/2021.06.11.448089

**Authors:** Hila Barzilai-Tutsch, Valerie Morin, Gauthier Toulouse, Stephen Firth, Christophe Marcelle, Olivier Serralbo

## Abstract

The Wnt/β-catenin signaling pathway is highly conserved throughout evolution and it plays crucial roles in several developmental and pathological processes. Wnt ligands can act at a considerable distance from their sources and it is therefore necessary to examine not only the Wnt-producing but also the Wnt-receiving cells and tissues to fully appreciate the many functions of this pathway. To monitor Wnt activity, multiple tools have been designed which consist of multimerized Wnt signaling response elements (TCF/LEF binding sites) driving the expression of fluorescent reporter proteins (e.g. GFP, RFP) or of LacZ. The high stability of those reporters leads to a considerable accumulation in cells activating the pathway, thereby making them easily detectable. However, this makes them unsuitable to follow temporal changes of the pathway’s activity during dynamic biological events. Even though fluorescent transcriptional reporters can be destabilized to shorten their half-lives, this dramatically reduces signal intensities, particularly when applied *in vivo.* To alleviate these issues, we developed two transgenic quail lines in which high copy number (12x or 16x) of the TCF/LEF binding sites drive the expression of destabilized GFP variants. Translational enhancer sequences derived from viral mRNAs were used to increase signal intensity and specificity. This resulted in transgenic lines efficient for the characterisation of TCF/β-catenin transcriptional dynamic activities during embryogenesis, including using *in vivo* imaging. Our analyses demonstrate the use of this transcriptional reporter to unveil novel aspects of Wnt signaling, thus opening new routes of investigation into the role of this pathway during amniote embryonic development.

## INTRODUCTION

The Wnt/β-catenin signaling pathway is a highly conserved pathway, which appeared early in phylogenesis and is common to all metazoan life forms. This pathway plays crucial roles during the entire lifespan of all organisms, from early embryogenesis to homeostasis in the adult. Wnt ligands bind to the transmembranal Frizzled receptor family and along with members of the LRP transmembanal coreceptor family, they mediate a large array of cell responses including cell fate specification, polarization, migration and mitogenic stimulation (Clevers, 2006; Clevers and Nusse, 2012; Logan and Nusse, 2004). The first and best characterized (and referred to as ‘‘canonical”) cellular response to Wnt is the inhibition of the β-catenin destruction complex, with the consequence of an increase of the β-catenin pool in the cytoplasm, ultimately leading to its translocation into the nucleus. There, it partners with members of the TCF/LEF family of transcription factors to activate various Wnt target genes, in a context-dependent manner.

To monitor the activity of the Wnt canonical pathway *in vitro,* transcription-based reporter systems were created by combining the DNA binding sites of TCF/LEF upstream of a minimal promoter and a reporter gene. The first reporter of Wnt/β-catenin signaling, TOPFlash, was used *in vitro* and it contained three TCF/LEF response elements upstream of a basal *c-fos* promoter driving the expression of the luciferase gene (Korinek et al., 1997; see also the Wnt Homepage for a complete list of references http://web.stanford.edu/group/nusselab/cgi-bin/wnt/). More sensitive TOPflash reporters were generated in *Drosophila* and zebrafish by increasing the number of TCF/LEF sites to 8, 12 and 16 (DasGupta et al., 2005; Veeman et al., 2003). In an attempt to detect Wnt signaling activity in mouse, three TCF/LEF binding sites were associated to LacZ and used to generate the TOPGAL mouse line (DasGupta and Fuchs, 1999), thus allowing the first analysis of Wnt responses in a vertebrate embryo. A significant increase in sensitivity was achieved by expanding the number of TCF/LEF binding sites to seven (BAT-gal mouse line; Maretto et al., 2003). Fluorescent reporter proteins (GFP or RFP and their variants) were also used in mouse and zebrafish (Ferrer-Vaquer et al., 2010; Moro et al., 2012). The main advantage of using either the β-galactosidase system or fluorescent proteins as reporters is their high stability: β-galactosidase half-life is reported to be up to 48 hours (Egan et al., 2013); that of GFP and RFP is about 24 hours (Corish and Tyler-Smith, 1999), and fusion of GFP to an H2B nuclear localization signal (as described in Ferrer-Vaquer et al., 2010) further stabilizes the fluorescent label (Foudi et al., 2009). Such high stabilities lead to a considerable accumulation of reporter proteins in cells activating the pathway, thus facilitating their detection. However, significant drawbacks are an important lag-time between the activation of the pathway and the detection of the reporter and, conversely, the detection of signals in tissues where Wnt activity may have already ceased. This makes stable reporters largely unsuitable to detect rapid spatiotemporal changes in a pathway activity. Destabilized fluorescent reporters have been designed to alleviate this problem, however shortening their half-life leads to dramatic fluorescence signal losses: for instance, d2GFP (half-life 2 hours), is 90% less fluorescent than its native GFP counterpart (He et al., 2019). Combining four TCF/LEF binding sites with a destabilized fluorescent reporter (d2EGFP, 2 hrs half-life, Clontech) in Zebrafish generated a transgenic line in which only intense activities of the pathway were detected through native fluorescence (Dorsky et al., 2002), thus requiring the more sensitive technique of in situ hybridization to detect lower Wnt signaling activities in this line. Increasing the number of TCF/LEF binding sites to six (upstream of a minimal promoter, miniP, and d2EGFP) generated a fish line with four insertion sites, in which many of the known Wnt/β-catenin signaling-active sites were detected by native fluorescence, including through live imaging (Shimizu et al., 2012).

Alternative reporter lines were also created by utilizing Wnt transcriptional targets (e.g. Axin2 or LGR5) to generate transgenic or knock-in mouse lines (Barker et al., 2007; de Roo et al., 2017; Lustig et al., 2001; Van Amerongen et al., 2012; van de Moosdijk et al., 2020). While those lines have been useful to characterize the targets’ response to Wnt, they only partially cover all activities of the Wnt/β-catenin pathway, and they are based on stable reporters, unsuitable to study dynamic signaling.

Importantly, recent evidence suggests that mobilization of β-catenin from the cell membrane pool can also trigger the activation of TCF/LEF reporters in a WNT ligand-independent manner (Sieiro et al., 2016; Lau et al., 2015). This indicates that, even though TCF/LEF-based reporters faithfully reflect TCF/β-catenin transcriptional activity, it may not all be due to canonical Wnt signaling.

While strategies described above are mainly based on enhancing transcriptional activity of TCF/β-catenin reporters, very little has been done to reinforce their translational efficiency. Sequence elements in the 5’ and 3’ untranslated regions of mRNAs play crucial roles in translation and well characterized elements derived from plant and viruses have been successfully used in heterologous systems (cell culture and *Drosophila)* to considerably increase reporter protein yields (He et al., 2019; Pfeiffer et al., 2012).

Here, we generated two novel transgenic quail lines carrying TCF/LEF-responsive elements, using the technology we recently developed (Serralbo et al., 2020). In the first (named 12xTFd2GFP), we have increased the number of TCF/LEF repeats to twelve, upstream of a cytoplasmic, destabilized EGFP (d2EGFP). We had previously used this construct for electroporation of chicken embryonic tissues and shown that it is more sensitive to the activity of Wnt signaling than existing reporters containing three or eight TCF/LEF repeats (Rios et al., 2010; Sieiro et al., 2016).

In the second line (named 16xTF-VNP), we increased transcriptional activity further, using sixteen TCF/LEF repeats. This was combined with translational enhancers to drive the expression of a nuclear, destabilized, fast maturing Venus (Nagai et al., 2002) as reporter.

This resulted in two transgenic lines in which the characterisation of the TCF/β-catenin transcriptional activity during embryogenesis is readily observed, including using *in vivo* imaging. Particularly remarkable is the 16xTF-VNP line, where intense, yet dynamic, reporter activity unveils unexpected features of TCF/β-catenin-responding cells and tissues.

This study underlines the importance of developing novel strategies to generate reporters efficient for monitoring of the spatiotemporal dynamics of signaling pathways and it opens new routes of investigation for correlating TCF/β-catenin transcriptional activities with unique cell behaviors during amniote embryonic development.

## RESULTS AND DISCUSSION

### Generation of the 12xTFd2GFP and 16xTF-VNP transgenic quail lines

To generate the 12xTFd2GFP line (TgT2(12TCF/LEF:d2EGFP)), we modified a TCF/β-catenin transcriptional reporter we previously generated, which was intended for in vivo electroporation in chicken embryos (Rios et al., 2010; Sieiro et al., 2016) by inserting Tol2 (T2) transposable elements 5’ and 3’ of the construct, thus allowing its stable integration into the quail genome (Fig. 1A). We used the direct injection technique as described in (Serralbo et al., 2020; Tyack et al., 2013) to transfect *in vivo* the blood-circulating primordial germ cells (PGCs). Fifty wild-type embryos at stage HH16 (E2.5) were injected in the dorsal aorta with a mix of lipofectamine 2000, the 12xTF-d2EGFP plasmid and a pCAG-Transposase construct. Four founders were selected, of which one male was used as founder. It was mated with wild-type females and their embryos were used for the experiments. The transmission of the transgene to the offspring presented a Mendelian distribution, suggesting a single insertion.

**Fig. 1.**
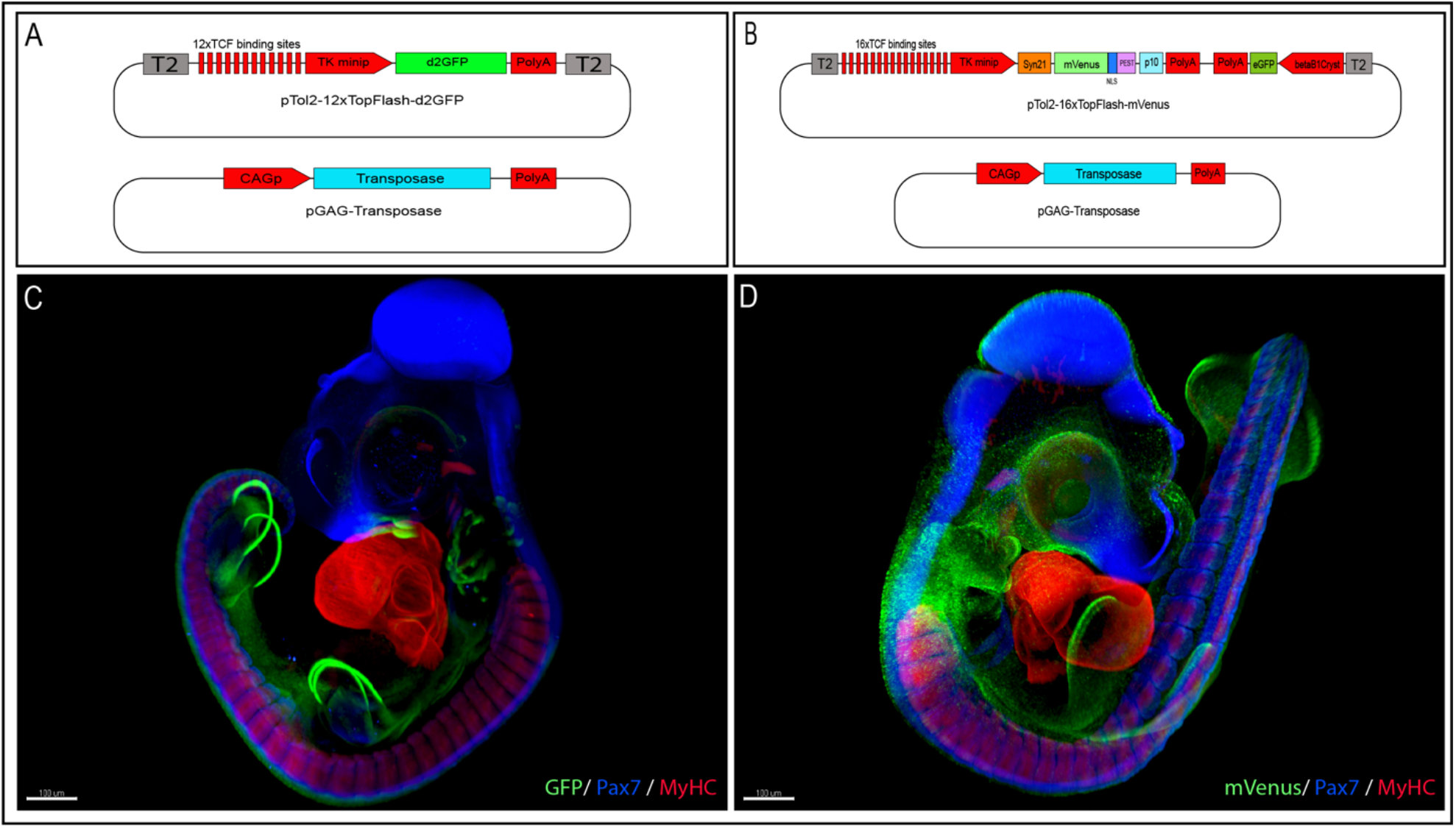
Generation of the 12xTFd2GFP and the 16xTF-VNP transgenic Japanese quail lines. **(A)** Vector used to generate the 12xTFd2GFP line and **(B)** the 16xTF-VNP line. **(C)** An E3.5 12xTFd2GFP embryo and **(D)** an E3 16xTF-VNP embryo cleared with the 3DISCO method, showing an overview of the TCF/β-catenin reporter activities in these lines. Embryos stained for Pax7 (Blue), MyHC (Red) and GFP or mVenus (Green). Scale bar 100 μm.

To further improve the TCF/β-catenin reporter sensitivity, we generated the 16xTF-VNP line (TgT2(16TCF/LEF:Syn21-Venus-NLS-PEST-p10, Gga.CRYBB1:GFP)). For this, we synthesized and cloned 16 TCF/LEF repeats upstream of the TK minimal promoter, followed by the Syn21 sequence (Pfeiffer et al., 2012), directly abutting the ATG initiation codon of Venus (Fig. 1B). Syn21 consists of an AT-rich 43-bp consensus translation initiation sequences made of elements derived from *Drosophila* and from the *Malacosoma neustria* nucleopolyhedrovirus polyhedrin gene. We chose a nuclear, destabilized form of the EYFP variant Venus as reporter, as it displays a 156% increase in relative brightness compared to EGFP (Nagoshi et al., 2004). It was followed by the p10 sequence, which is a 606 bp terminator sequence from the *Autographa californica* nuclear polyhedrosis baculovirus (Pfeiffer et al., 2012). A CrystallGFP selection mini-gene, consisting of the promoter of the βB1crystallin gene upstream of EGFP was added to the construct to ease the selection of transgenic birds at hatching (Serralbo et al., 2020). Finally, Tol2 sites were added for stable integration into the quail genome.

To validate the efficiency of this construct, we electroporated one half of the neural tube of 15HH (E2) chicken embryos with a DNA mix containing the 16xTF-VNP construct and a TagBFP protein driven by the CAG ubiquitous promoter as a marker of electroporated cells. Twenty-four hours after electroporation, while one half of the neural tube and migrating neural crest cells displayed strong expression of the electroporation marker (TagBFP), weak expression of 16xTF-VNP was observed in the dorsal neural tube (NT) while stronger expression of the reporter was seen in a portion of the neural crest cell population (NC; Sup. Fig. 1A-D). To compare the sensitivity of the 16xTF-VNP construct to that of the 12xTF-d2EGFP; we co-electroporated the neural tube of E2 chicken embryos with either the 12xTF-d2EGFP or the 16xTF-VNP together with CAG-TagBFP, as internal control (Sup. Fig. 1E-G and H-J, respectively). Twenty-four hours after electroporation, we calculated the percentage of the d2EGFP or mVenus positive NC cells out of the total TagBFP cells and we observed a 2.5-fold increase in the percentage of the mVenus-positive cells compared to the d2EGFP cells (Sup. Fig.1N). To verify that the accumulation of TCF/LEF binding sites did not lead to higher basal level of activation, we inhibited Wnt signaling in the presence of the reporter. The dorsal NT of E2 chicken embryos was coelectroporated with the 16xTF-VNP and the CAG-TagBFP together with a dominant-negative LEF1 (DN-LEF1; Sup. Fig. 1K-M). Twenty-four hours post electroporation we observed a 5-fold reduction in the percentage of mVenus-positive cells in the embryos co-electroporated with the DN-LEF1 as opposed to the controls (Sup. Fig. 1O). Altogether, this suggests that the 16xTF-VNP construct is a reliable and sensitive reporter of TCF/β-catenin transcriptional activity. Furthermore, as we had shown that the 12xTF-d2EGFP response was more intense than existing reporters (Rios et al., 2010), it is likely that the 16xTF-VNP represents the most sensitive reporter to date to monitor the dynamics of TCF/β-catenin transcriptional activity.

We used the same technique as above to generate transgenic quails with this reporter. One female was selected as the transgenic founder using the CrystallGFP marker and crossed with a WT male to expand the transgenic line. The transmission of the transgene to the offspring presented a Mendelian distribution, suggesting a single insertion.

### TCF/β-catenin transcriptional activities in early embryos

To characterize TCF/β-catenin transcriptional activity during embryonic development, we analyzed and compared the activity of the two reporters in the two lines we generated, in whole mount preparations of immunostained embryos, on immunostained sections and in live tissues at different developmental stages. Whole mount preparations of HH21 (E3.5; 12xTF-d2GFP) and HH20 (E3; 16xTF-VNP) transgenic quail embryos immunostained for GFP, Pax7 and Myosin Heavy Chain (MyHC), clarified by the ‘3DISCO’ technique (Belle et al., 2017) and imaged using selective/single plane illumination microscopy (SPIM, LaVision BioTec UltraMicroscope II), give an overview of the sites of high reporter activities. It shows conspicuous reporter activity in the Apical Ectodermal Ridge (AER), the pharyngeal arches (particularly the maxillary), the somites, etc. (Fig. 1C,D; Sup. movies 1 and 2). This initial examination also indicated that the 16xTF-VNP reporter line is more sensitive to TCF/β-catenin transcriptional activity than the 12xTF-d2GFP, as it shows reporter activity in places that are not detected in the 12xTF-d2GFP line (e.g. the cephalic neural crest; Fig 1D).

To further characterize TCF/β-catenin activity during embryonic development, we performed immunostaining on sections and on whole mount embryos, coupled with classical confocal microscopy, which are more sensitive techniques than the SPIM. Transverse sections of E3 12xTF-d2GFP and 16xTF-VNP embryos stained for GFP, Pax7 and MyHC confirmed that TCF/β-catenin transcriptional activity was generally stronger in the 16xTF-VNP than in the 12xTF-d2GFP. For instance, reporter activity was observed throughout the dermomyotome (stronger in the medial and lateral border, DML and VLL, respectively) in the 16xTF-VNP, while it was visible only in the DML and VLL in the 12xTF-d2GFP. In both reporter lines, significant reporter expression was present in mesenchymal cells within the limb bud, and strong activity was observed in the AER (Fig 2 A-D).

**Fig. 2.**
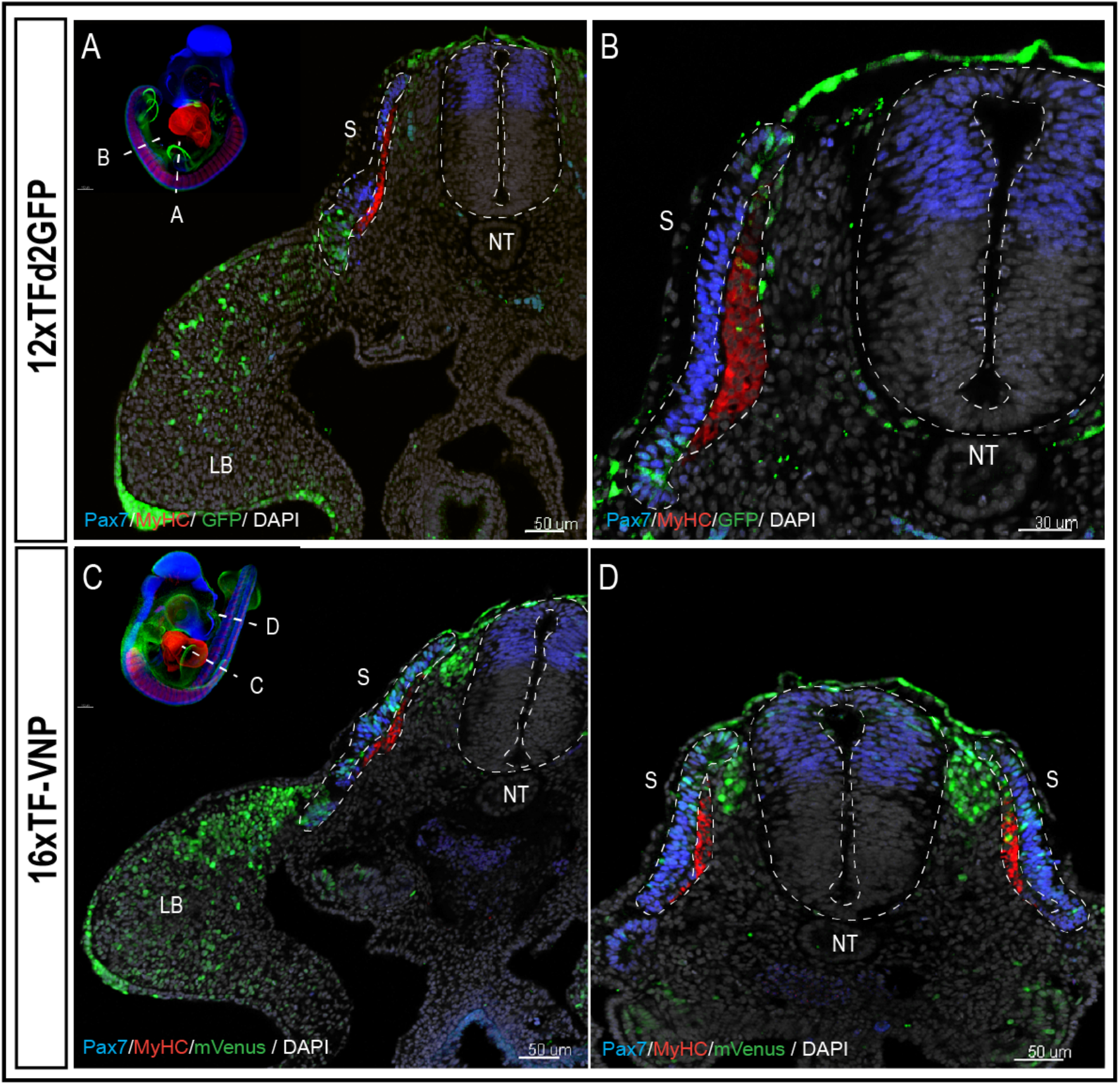
The 16xTF-VNP reporter is a sensitive reporter of the TCF/β-catenin signaling activity. Transverse sections at the levels of the front limb (**A** and **C**) and the trunk **(C** and **D**) of E3 12xTFdsGFP **(A-B)** and 16xTF-VNP **(C-D)** transgenic embryos stained for GFP or mVenus (green), Pax7 (Blue), MyHC (red) and DAPI (grey). Inserts show the levels at which the sections were made. Scale bar 50 μm. S = somite, NT = neural tube, LB = limb bud.

We made surprising observations in the neural tube and neural crest (NC). We observed that the reporter was inactive in the dorsal neural tube in the 12xTF-d2GFP (Fig 2 A, B and Sup Fig 2 A-J) and weakly active in the more sensitive 16xTF-VNP line (Fig 2 C,D, and Sup. Fig 2 K-T). In contrast, migrating NC strongly upregulated the reporter as they left the neural tube *en route* to their sites of differentiation (this is particularly visible in the 16xTF-VNP; Fig 2 C,D, and Sup. Fig 3 A-D). These patterns of the reporters’ activities are unexpected, in regard to the published expression patterns of Wnt mRNAs and proteins in those tissues. Previous works have shown that Wnt1 and Wnt3a mRNA transcripts are present in the roof plate (RP) of the neural tube in mouse and chicken early embryos (Hollyday et al., 1995; Marcelle et al., 1997), while their transcripts are absent from migrating NC. Wnt expression is maintained in the dorsal neural tube at least as long as neural crest cells are generated from the RP (Sup Fig 4 A-D). We recently uncovered that Wnt1 protein (Wnt3a was not tested), under the control of BMP3 and 7 in the dorsal neural tube (Marcelle et al., 1997) and detected by immuno-histochemistry in RP cells, is loaded onto NC as they initiate their migration to be delivered at a distance to somites where it serves to regulate myotome organization (Serralbo and Marcelle, 2014). The observation we made here brings another level of understanding of the roles of Wnt in those tissues, since it indicates that at the time of observation, epithelial cells of the dorsal neural tube and the RP poorly respond to Wnt, while the epithelial-mesenchyme transition necessary for NC migration triggered a strong increase in the reporter’s response in those cells. As NC proceeded along their dorso-ventral migration path, the reporter activity rapidly diminished, generating a gradient of TCF/β-catenin transcriptional activity likely due to the exhaustion of the pool of Wnt ligand initially loaded on the NC cell surface (Fig 2 C,D, and Sup. Fig 3 A-D). The functions of Wnt signaling in the CNS and the NC are unclear. Loss and gain-of-function of Wnt1 and/or 3a in mouse, xenopus and chicken have led to the premise that these molecules play important functions in the proliferation and/or differentiation of neuronal precursors within the CNS, opposing a Shh gradient from the notochord and floor plate (Dickinson et al., 1994; Ikeya et al., 1997; Mcmahon et al., 1992; Saint-Jeannet et al., 1997; Alvarez-Medina et al., 2008; Dessaud et al., 2008). However, patterning defects in Wnt1 or compound Wnt1/Wnt 3a mutants were only observed in the brain, while the spinal cord was unaffected (Ikeya et al., 1997; Mcmahon et al., 1992). It is in fact possible that the function of Wnts in the CNS varies in time and place during embryogenesis.

Loss of Wnt-1 and Wnt-3a function in mouse resulted in a large deficit in all neural crest derivatives (e.g. in cranial and dorsal root ganglia), that suggested a role in the neural crest cell proliferation (Ikeya et al., 1997). Surprisingly, both the inhibition or the activation of Wnt signaling in neural crest cells were reported to promote neuronal fates at the expense of other derivatives (Dorsky et al., 1998; Lee et al., 2004). The observations made in the quail lines we generated cannot differentiate between these contradictory hypotheses, but they clarify the time window and the place when Wnt signaling is likely to be active in these processes (i.e. the early migratory phases) and may explain some of the changes observed in the transcriptional signature of neural crest cells along their migratory routes (Azambuja et al., 2021; Simões-Costa and Bronner, 2015; Morrison et al. 2017).

A similar gradient of reporter activity was observed in somites since it was active in the DML and the transition zone and inactive in fully elongated myocytes of the myotome (Fig. 3 A-D). As previously described, high and low TCF/β-catenin transcriptional activities were observed within DML cells, the strongly labelled ones corresponding to DML cells engaging into the myogenic program (see also below; Sieiro et al., 2016).

**Fig. 3.**
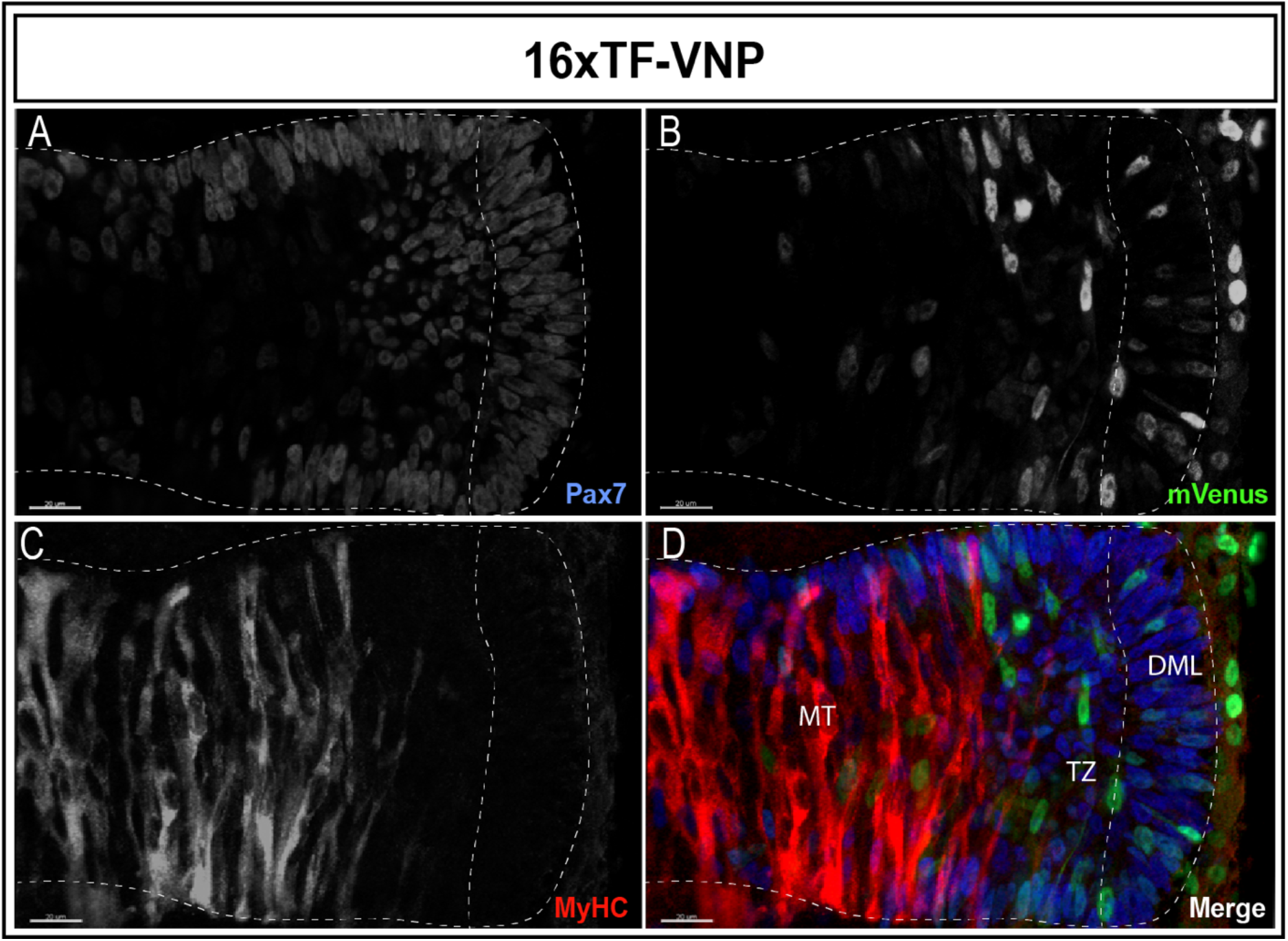
A gradient of dynamic TCF/β-catenin signaling activity in somites. An optical section of an E2.5 embryo somite stained for Pax7 (**A,D**; blue) mVenus (**B,D**; green), and MyHC (**C,D**; red) reveals high and low reporter expression levels in the DML and strong expression in elongating myocytes located in the transition zone (TZ). The reporter levels are rapidly decreased in the myotome. Scale bar 20 μm. DML = dorso-medial lip, MT = myotome.

Thus, the use of destabilized fluorescent reporters to generate these quail lines allows the detection of dynamic changes (increase and decrease) in reporter activity throughout embryonic development that could not be appreciated with stable reporters, such as the BAT-gal mouse line (Maretto et al., 2003).

### TCF/β-catenin activity during late organogenesis

We also analyzed the activity of the reporters at 9 days of development (HH35; Fig. 4). Tissues that displayed strong TCF/β-catenin transcriptional activity include the egg tooth (Fig. 4A,F), the liver (Fig. 4B,F), the feather buds (Fig. 4C), the embryonic vertebrae (Fig. 4D), as well as the limb bone growth zones (Fig. 4E). While the significance of the conspicuous reporter activity in egg tooth formation is unknown, Wnt functions in liver, feather follicles and bone formation during organogenesis are well documented. Wnt/β-catenin was shown to promote hepatocyte proliferation in mice (Perugorria et al., 2019), to initiate the formation of hair and feather follicle placodes in mouse and chicken (DasGupta and Fuchs, 1999; Fuchs, 2016; Noramly et al., 1999; Olivera-Martinez et al., 2001) and to favor osteoblast differentiation over chondrocyte differentiation, thereby determining whether mesenchymal progenitors become osteoblasts or chondrocytes (Baron et al., 2006; Day et al., 2005).

**Fig. 4.**
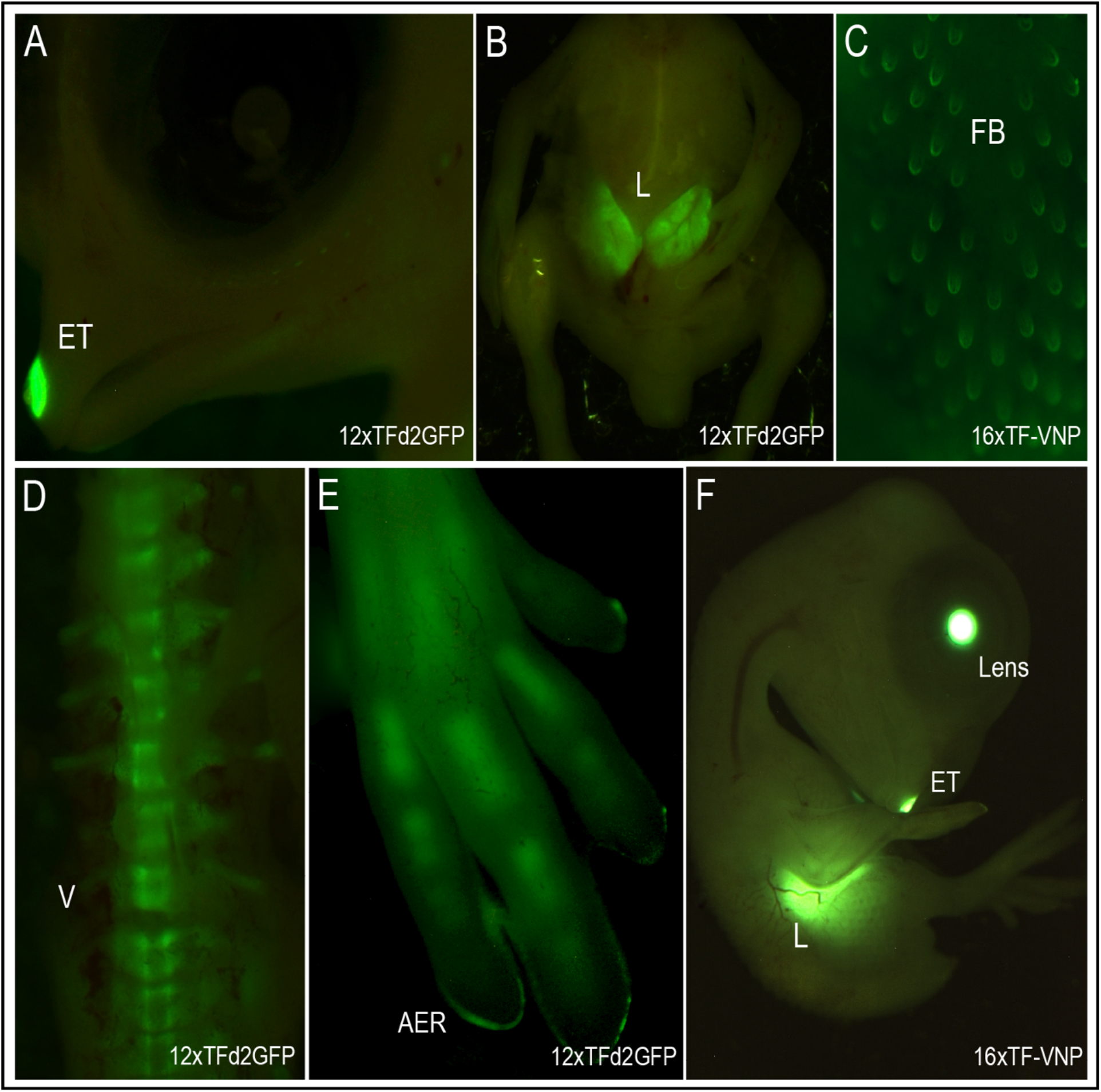
TCF/β-catenin reporter activity in E9 transgenic embryos. Native TCF/β-catenin signalling reporter in E9 12xTFd2GFP **(A,B,D,E)** and E8 16xTF-VNP embryos **(C,F)**. The TCF/β-catenin reporter is strongly expressed in the egg tooth, liver and AER of 12xTFd2GFP (**A,B,E** respectively) and 16xTF-VNP (**F**) embryos as well as in the bones of 12xTFd2GFP embryos (**D,E**). The reporters are also expressed in feather follicles (**C**). In the 16xTF-VNP embryo the CrystallGFP selection marker is strongly visible in the lens (**F**). ET = egg tooth, L = liver, FB = feather bud, V = vertebrae, AER = apical ectodermal ridge.

### Dynamic TCF/β-catenin transcriptional activity in live tissues

TCF/β-catenin spatiotemporal activity is highly dynamic, both on a cellular level and throughout development, a feature that has been difficult to study due to lack of reliable destabilised reporters. By using the 12xTFd2GFP and the 16xTF-VNP transgenic lines, we followed the dynamics of TCF/β-catenin transcriptional activity *in vivo.* Observation of early somites showed rapid and strong activation of the 12xTFd2GFP reporter activity in single cells located in the medial portion of the somite (named the DML (Sup. Video 3, arrowheads; see also Sup Fig 5). This is coherent with our previous studies, which showed that single epithelial cells within the DML, receiving Delta signals from incoming migrating neural crest cells, respond by activating the myogenic program through a NOTCH/β-catenin-dependent/Wnt-independent signaling module (Rios et al., 2011; Sieiro et al., 2016). The in vivo analysis presented here suggest that the entry of DML cells into myogenesis can be monitored through an increase in 12xTFd2GFP activity and their behavior followed live using the quail lines we generated. An interesting finding of this study was the intense TCF/β-catenin-response in the fore- and hindlimb Apical Ectodermal Ridges (AER). The AER remained strongly labeled throughout limb growth (up to E9, where it was still detected at the finger tips, Fig 4E). Interestingly, the reporter^+^ cells were found scattered in a wide region of early limb ectoderm (Sup. Video 4), intermingled with reporter-negative cells. As the limbs grew, reporter^+^ cells migrated towards and coalesced to form the AER, suggesting that they constitute a population of AER progenitors. The AER is crucial to limb formation, which serves as a signaling center that regulates dorso-ventral patterning of the limb and its proximo-distal growth (Fernandez-Teran and Ros, 2008; Zeller et al., 2009). Interestingly, lineage analyses performed in chicken embryos had shown that the AER is derived from cells located on a wide region of early ectoderm (Altabef et al., 1997; Michaud et al., 1997) and suggested that AER progenitors were intermingled with non-ridge progenitor, an observation coherent with our finding. Wnt3a in the chicken embryo follows a similar pattern to that of the reporter, widely expressed throughout the dorsal ectoderm in early developing limb buds, and later condensing to the AER region (Fernandez-Teran and Ros, 2008; Zeller et al., 2009; Kengaku et al., 1998). While it is possible that AER progenitors (recognized by their expression of the TCF/β-catenin reporter) are specified and/or respond to the Wnt3a signal, a recent study suggested that mechanical tensions from the underlying growing limb mesenchyme participate in the activation of TCF/β-catenin response (and therefore in the specification of AER progenitors) in the overlying ectoderm in a Wnt-independent manner (Lau et al., 2015). In this context, it will be interesting to determine whether Wnt3a acts as a directional cue in the migration of AER progenitors toward the AER anlage.

In summary, we have generated two transgenic quail lines suitable for the study of the dynamic behaviour of TCF/β-catenin signaling, with the 16xTF-VNP likely being the most sensitive transgenic line to date. Both lines allow overcoming previous limitations in the study of the Wnt/TCF/ β-catenin signaling, particularly in vivo and their availability opens new routes of investigation into dynamic signaling activity of this pathway throughout development.

## MATERIALS AND METHODS

### Generating transgenic quail by direct injection

The direct injection technique is performed as described in (Tyack et al., 2013; Serralbo et al., 2020). The injection mix contained 0.6ug of Tol2 plasmid, 1.2ug of CAG-Transposase plasmid, 3ul of lipofectamin 2000 CD in 90 ul of Optipro. 1ul of injection mix was injected in the dorsal aorta of 2.5-day-old embryos. After the injection, eggs were sealed and incubated until hatching. Hatchlings were grown for 6 weeks until they reach sexual maturity. Semen from males was collected using a female teaser and the massage technique as described in (Chełmońska et al., 2008). The genomic DNA from semen was extracted and PCR was performed to test for the presence of the transgene in semen. Males showing a positive band in semen DNA were crossed with wild type females. Offspring were selected directly after hatching by PCR genotyping 5 days after hatching by plucking a feather. For the 16xTF-VNP birds, the CrystallGFP expression was also used for easy screen of transgenic birds.

### Whole-mount and sections immunochemistry and confocal analyses

Embryos were dissected under a Leica fluorescent stereomicroscope and fixed for 1h in 4% formaldehyde at RT. For sections preparations, embryos were embedded in 15% sucrose/7.5% gelatine/PBS solution and sectioned with Leica cryostat at 20μm. Antibody staining was performed as described in (Serralbo and Marcelle, 2014). The following primary antibodies were used: anti-GFP chicken polyclonal (ab13970, Abcam;1/1000), anti-Pax7 IgGI mouse monoclonal (Hybridoma Bank; 1/10), Rabbit polyclonal anti-Sox9 (ab5535, Millipore; 1/1000), anti-Myosin heavy Chain IgGIIb (Hybridoma bank; 1/10), anti-HNK1 IgGM (Hybridoma bank; 1/10). Images of stained sections were acquired with a Leica SP5 upright confocal microscope and an UV-corrected HCX PL APO CS 40x / NA 1.25 Oil immersion objective (WD 0.1 mm), combined with tile scan acquisition. For whole-mount samples, a CARL ZEISS LSM 980 Airyscan 2 confocal microscope on inverted Axio Observer stand was used, with UV-IR corrected PL APO 40x/1.3 oil immersion objective (WD 0.20 mm).

Images of native reporter activity in E9 embryos were taken under a Leica 3D Fluorescent microscope 4x dry objective.

### 3DISCO clearing

Performed as described in (Belle et al., 2017). In short, embryos were dissected under a Leica fluorescent stereomicroscope and fixed for 1h in 4% formaldehyde at RT. The embryos were immunostained as described above. Following immunostaining, embryos were dehydrated by immersion in 50%, 70%, 80% and 100% of tetrahydrofurane (THF; in milli-Q water). After dehydration the embryos were rinsed in dichlormethane (DCM) and finally in dibenzyl ether (DBE) to match the refractive index of tissue and surrounding medium leading to transparent sample. Images were acquired on LaVision BioTec UltraMicroscope II based on upright Olympus MVX10 macro zoom microscope with MVPLAPO 2 XC objective (NA 0.50, WD 20 mm) with 2x optical zoom. Overall stack thickness 3 μm.

### Time-lapse imaging

For time-lapse imaging, embryos were incubated and cultured in custom-made egg incubator described in (Serralbo et al. 2020, QuailNet database http://quailnet.geneticsandbioinformatics.eu/). Embryos were imaged using Leica SP8 confocal microscope on upright DM6000 stand with HCX APO L 20x/1.00 water dipping (WD 2.00 mm). Z-stack images were taken every 15 min for 9h (Sup. video 1) and 7h (Sup video 2). Wider field of view time-lapse imaging were acquired every 10mn for 10h (Sup. Video 3) and 9h (Sup.video 4) using Leica Thunder Image Model Organism. Images stitched together using the Image J software with drift correction plugin.

## ACKNOWLEDGEMENTS

The authors thank Taryn Guinan from Leica Biosystem for her supervision on Thunder Image Model Organism stereo microscope. The authors acknowledge Monash Micro Imaging and in particular Oleksandr Chernyavskiy, Monash University, for the provision of instrumentation, training and technical support with the lightsheet Ultramicroscope microscope. The Australian Regenerative Medicine Institute is supported by grants from the State Government of Victoria and the Australian Government. HBT was supported by grants from Stem Cell Australia and the Agence Nationale de la Recherche. We thank the Faculty of Medicine and Health Science for their financial support.

## Notes

### Competing Interest Statement

The authors have declared no competing interest.

