## Supplementary Figures for "TopFlash transgenic quail reveals dynamic TCF/β-catenin signaling during avian embryonic development"

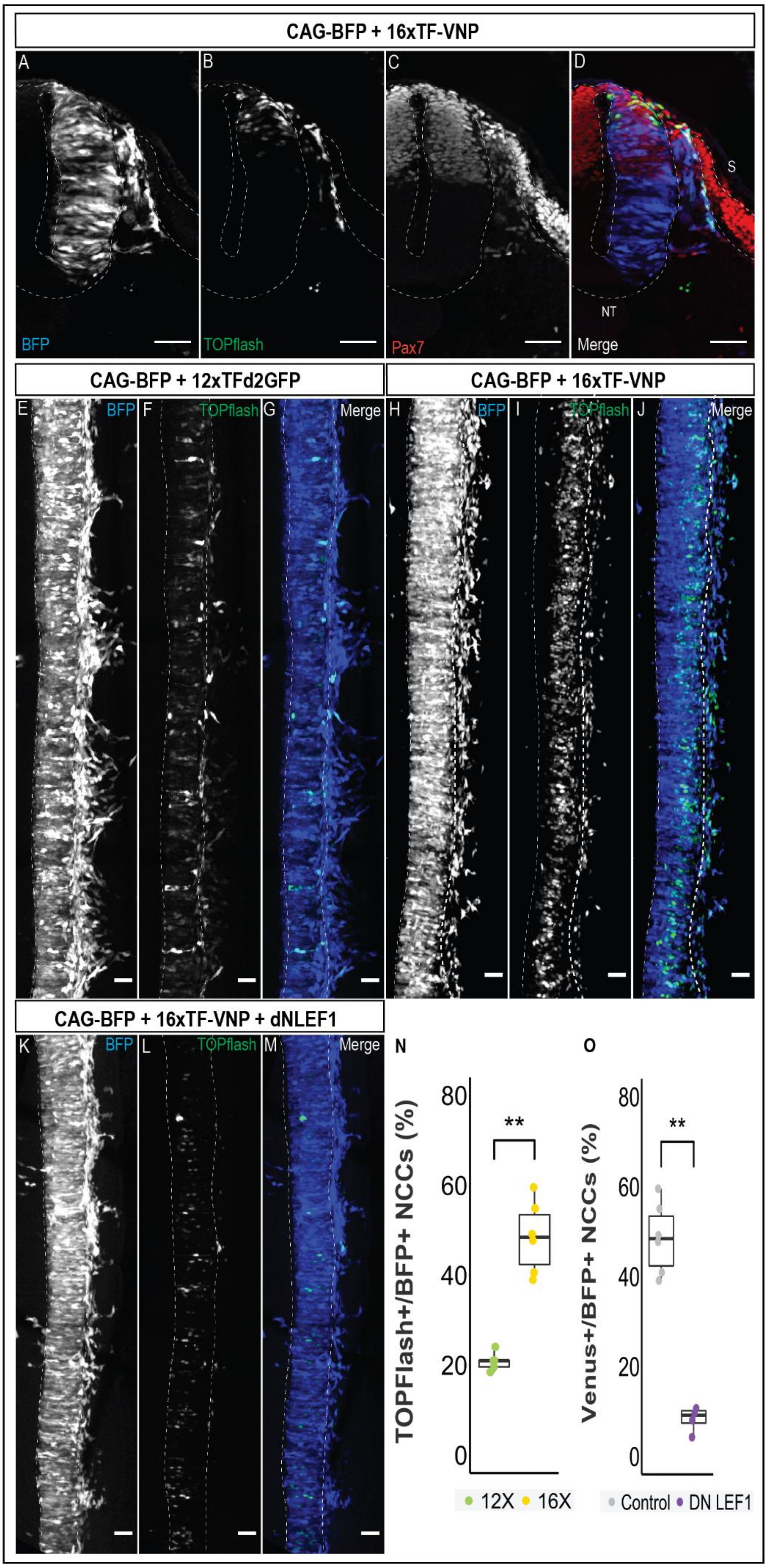

CAG-BFP + 16xTF-VNP
A
B
C
D
S
NT
BFP
TOPflash
Pax7
Merge
CAG-BFP + 12xTFd2GFP
CAG-BFP + 16xTF-VNP
E BFP
F TOPflash
G Merge
H BFP
I TOPflash
J Merge
CAG-BFP + 16xTF-VNP + dNLEF1
K BFP
L TOPflash
M Merge
N
O
TOPFlash+/BFP+ NCCs (%)
Venus+/BFP+ NCCs (%)
0
20
40
60
80
**
12X
16X
Control
DN LEF1

**Sup. Fig. 1 – Comparison of the 12xTFd2GFP and the 16xTF-VNP TCF/β-catenin signaling activities.**

**(A-D)** Transverse sections of an E2.5 chick embryo electroporated in the NT with a DNA mix containing a CAG-TagBFP construct as a marker for electroporated cells and the 16xTF-VNP construct. The sections were stained with BFP (blue), mVenus (green) and Pax7 (red) and demonstrate the reporter activation in the dorsal NT and migrating NC cells.

**(E-G)** Confocal stack of a dorsal view of an E2.5 chick embryo electroporated in the NT with a DNA mix containing a CAG-TagBFP construct as a marker for electroporated cells and the 12xTFd2GFP construct. BFP (blue), GFP (green).

**(H-J)** Confocal stack of a dorsal view of an E2.5 chick embryo electroporated in the NT with a DNA mix containing a CAG-TagBFP construct as a marker for electroporated cells and the 16xTF-VNP construct. BFP (blue), mVenus (green).

**(K-M)** Confocal stack of a dorsal view of an E2.5 chick embryo electroporated in the NT with a DNA mix containing a CAG-TagBFP construct as a marker for electroporated cells, the 16xTF-VNP construct and a DN-Lef construct demonstrate a downregulation of the TF-reporter compared to the control embryos **(A-D)**.

**(N)** Cell quantification shows a 2.5-fold increase in the number of cells electroporated with the 16xTF-VNP reporter compared to the 12xTFd2GFP (N=6 embryos per treatment).

**(O)** Cell quantification shows a 4-fold decrease in cells expressing the 16xTF-VNP reporter following a DN-Lef1 treatment. (N=6 embryos per treatment).

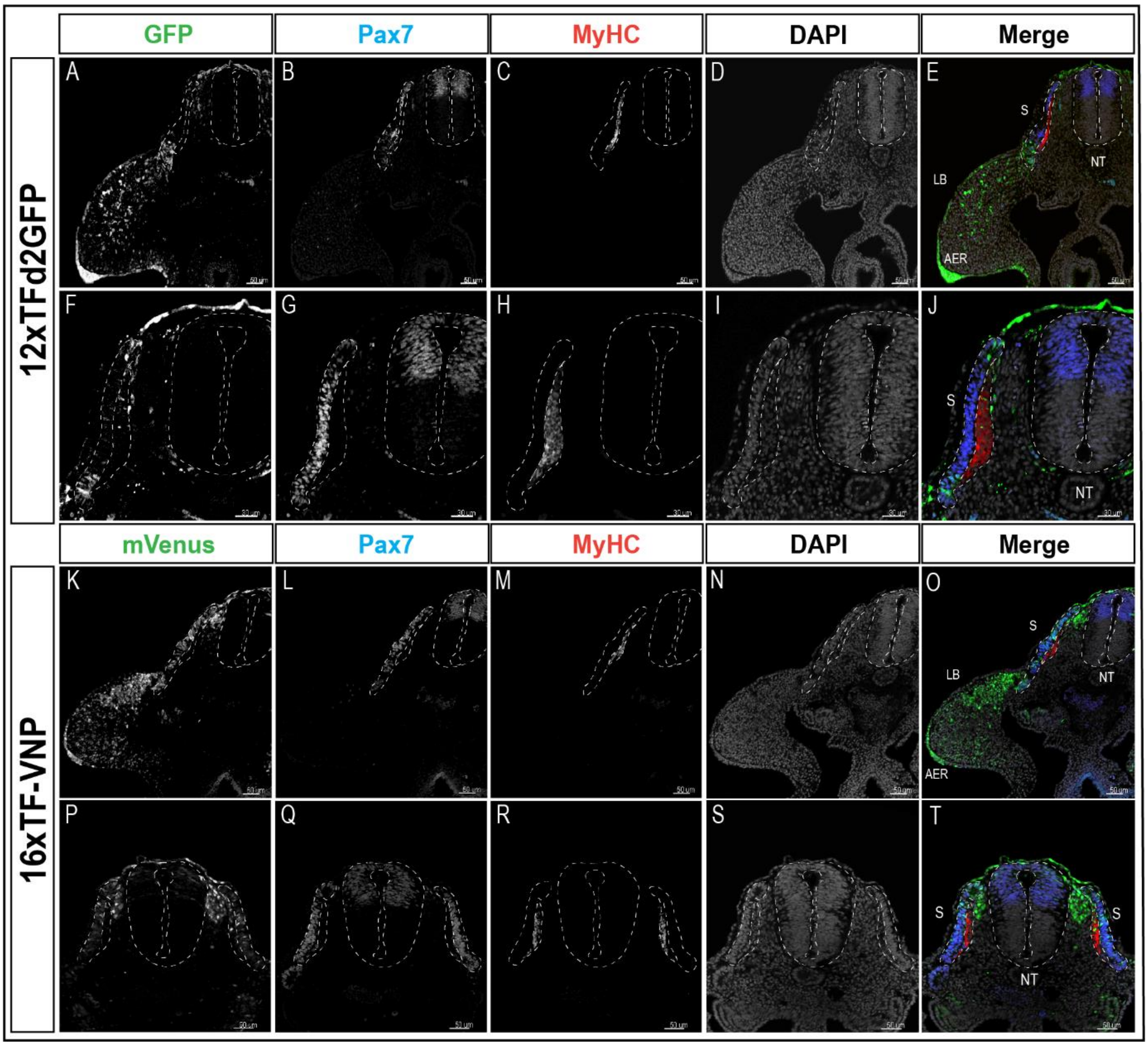


**Sup. Fig. 2 -**

Transverse sections at the levels of the front limb and the trunk of E3 12xTFd2GFP and 16xTF-VNP transgenic embryos stained for GFP or mVenus, respectively (green), Pax7 (Blue), MyHC (red) and DAPI (grey). The image details the different channels for Fig. 2. Scale bar 50 µm. S = somite, NT = neural tube, LB = limb bud, AER = apical ectodermal ridge.

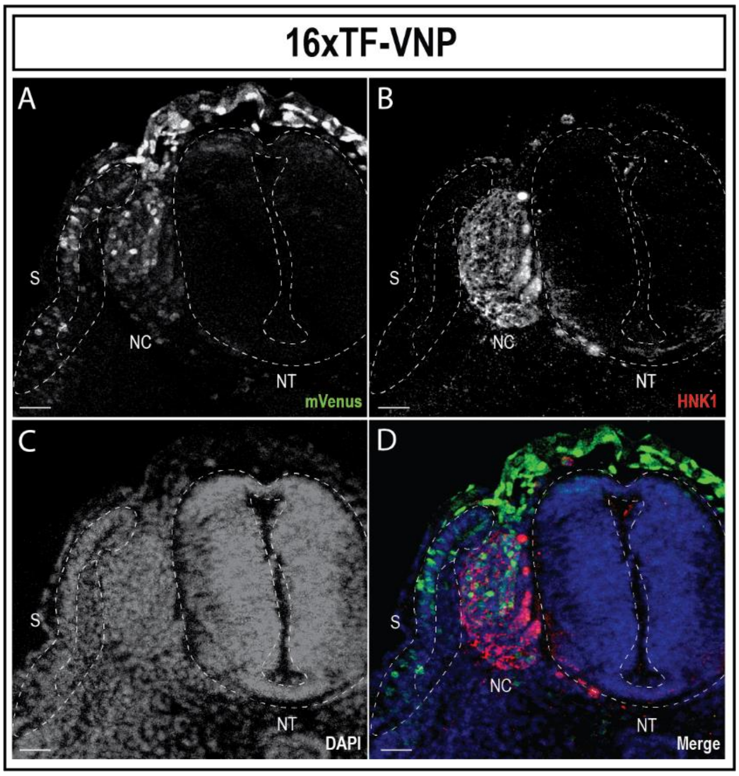


**Sup. Fig. 3 – TCF/β-catenin reporter dynamic activity in NC cells**Transverse sections of E3 (HH19) 16xTF-VNP stained for mVenus (green; **A,D**), HNK1 (red; **B,D**) and DAPI (blue; **C,D**). The TCF/β-catenin reporter is significantly upregulated in emerging NC cells and is rapidly decreased as the NC proceed along their dorso-ventral migration path. Scale bar = 50 µm. NT = neural tube, NC = neural crest cells, S = somite.

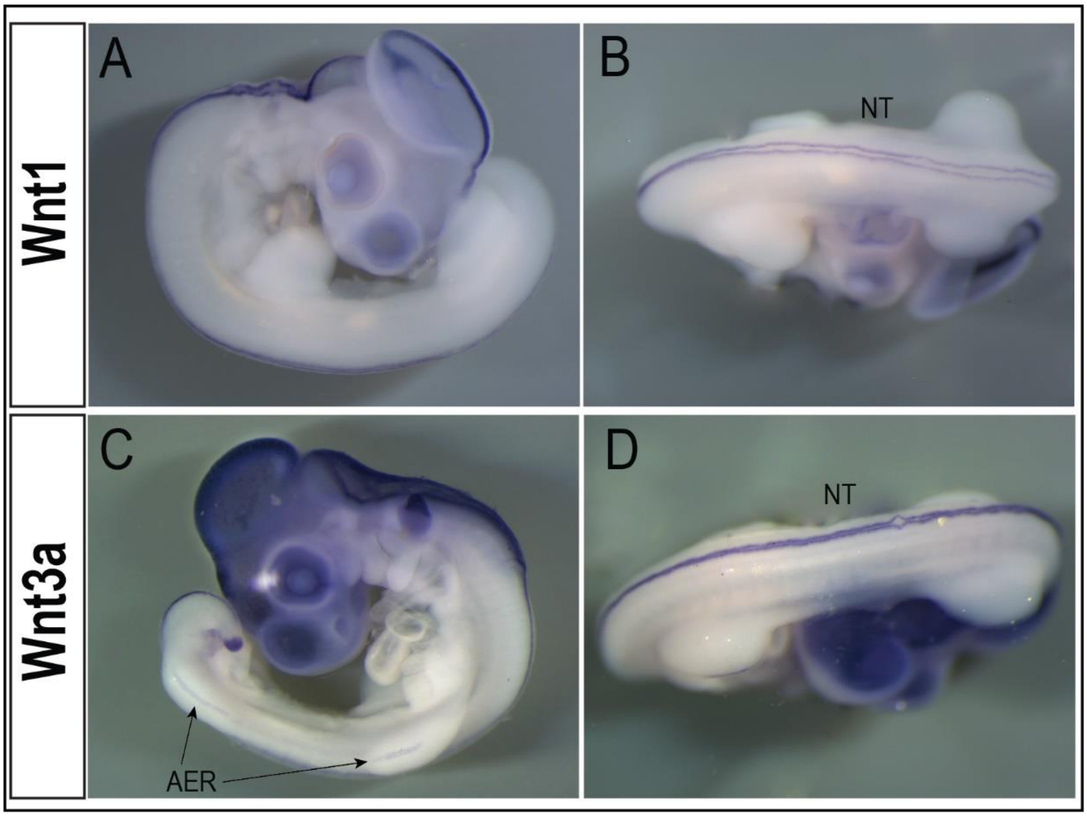


**Sup. Fig. 4 – In Situ Hybridization of Wnt3a and Wnt1 mRNA in quail embryos**

In Situ Hybridization of Wnt1 **(A,B)** and Wnt3a **(C,D)** in E3.5 quail embryos. Both Wnt1 and Wnt3a mRNA transcripts are presented in the NT (**A**, **B**,**D**). Wnt3a mRNA transcript is also present in the AER (**C**). NT = neural tube, AER = apical ectodermal ridge.

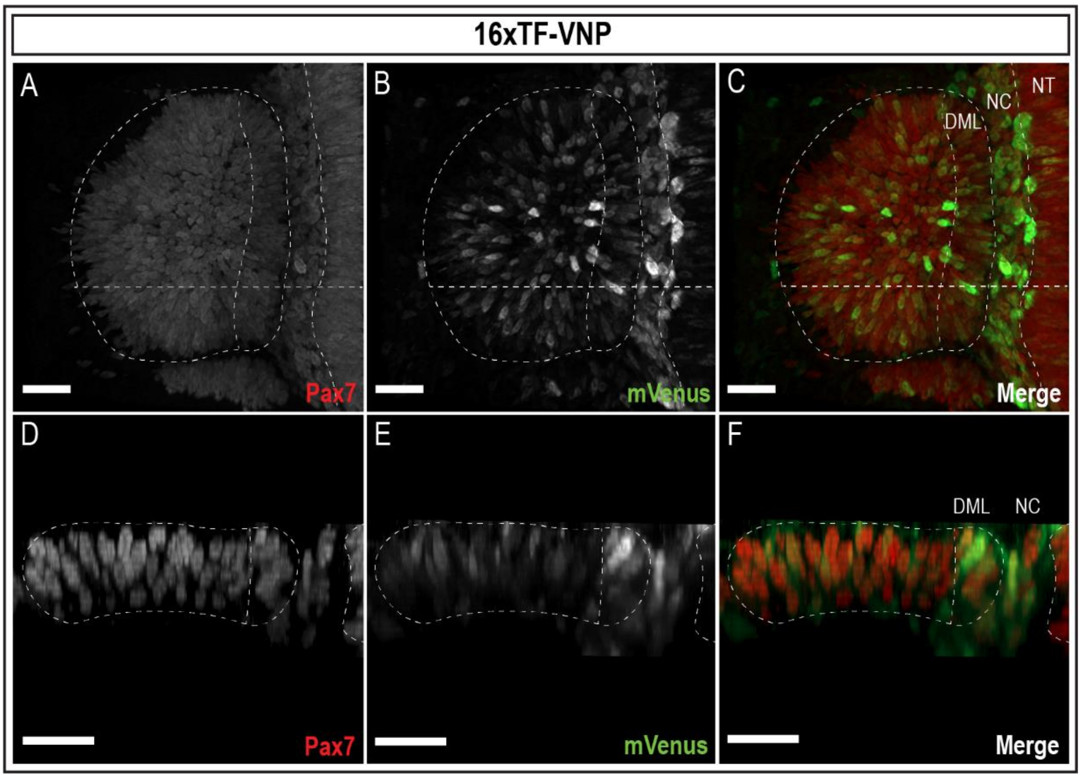


**Sup. Fig. 5 -**

3D reconstruction of a Z-stack of an E2.5 16xTF-VNP embryo somite shown in a dorsal **(A-C)** and transverse view **(D-F)** demonstrates reporter-positive migrating NC cells as well as different expression levels of the reporter across the somite and especially in the DML. Cells stained for mVenus (Green) and Pax7 (Red). Scale bar = 50 µm. NT = neural tube, NC = neural crest cells, DML = dorso-medial lip.

**Sup. Video 1 –**

A 3D reconstruction of E3.5 12xTFd2GFP embryo cleared with the 3DISCO method. The embryo is stained for GFP (green), Pax7 (blue) and MyHC (red).

**Sup. Video 2 –**

A 3D reconstruction of E3 16xTF-VNP embryo cleared with the 3DISCO method. The embryo is stained for GFP (green), Pax7 (blue) and MyHC (red).

**Sup. Video 3 –**

A 9h time-lapse confocal analysis of an E2.5 12xTFd2GFP embryo somite. The movie shows a single Z-plane of the somite (10um) and focuses on two cells in the dorsal DML which increase the TCF/β-catenin reporter activity (magenta arrowheads). We also show a cell division event in the caudal DML (white arrowheads).

**Sup. Video 4 –**

A 9h time-lapse movie of an E2 16xTF-VNP embryo showing the growing hind limb bud. Ectodermal cells strongly expressing the TCF/β-catenin reporter are seen as they migrate towards the AER region where they condensate.

The migrating NC and somitic cells are also highly visible.
